# Comprehensive longitudinal study of home-cage activity, including climbing, reveals new complex phenotypic profile in the N171-82Q HD mouse model with implications for refined preclinical studies

**DOI:** 10.1101/2023.01.19.524755

**Authors:** Rasneer S. Bains, Hamish Forrest, Rowland R. Sillito, J. Douglas Armstrong, Michelle Stewart, Patrick M. Nolan, Sara E. Wells

**Affiliations:** Mary Lyon Centre at Medical Research Council, Harwell, Oxfordshire, UK; Medical Research Council, Harwell Science Campus, UK; Actual Analytics Ltd., Edinburgh, UK; School of Informatics, University of Edinburgh, UK

**Author notes:** **Correspondence:** Rasneer S Bains.

**Keywords:** Automated, Neurodegeneration, Motor Function, Reproducible, Welfare, Huntington’s

## Abstract

Monitoring the activity of mice within their home cage is proving to be a powerful tool for revealing subtle and early-onset phenotypes in mouse models. Video tracking, in particular, lends itself to automated machine-learning technologies that have the potential to improve the manual annotations carried out by humans. This type of recording and analysis is particularly powerful in objective phenotyping, monitoring behaviors with no experimenter intervention. In this study, we focus on non-evoked voluntary behaviors, which do not require any contact with the animal or exposure to specialist equipment. We show that the monitoring of climbing on the wire cage lid of a standard individually ventilated cage (IVC) yields reproducible data reflecting complex phenotypes of individual mouse inbred strains and of a widely used mouse model of neurodegeneration. In addition, performing such measurements in the home-cage environment, over several 24-hour periods, allows for the collection of comprehensive behavioral and activity data, which reveals prolific sexual dimorphism and biphasic changes in locomotor activity. Here we present data from home-cage analysis, which reveals the complexity of unprovoked behavior in both wild-type and mutant mice. This has the potential to greatly enhance the characterization of mouse strains, detect early and subtle signs of disease and increase reproducibility in preclinical studies.

## 1 Introduction

Advances in the field of genetics mean that mouse models are increasingly sophisticated, and more closely than ever before model human disease (Mingrone et al., 2020). For diseases affecting neuromuscular systems, there is also an increasing range of assays and tests for mice that measure parameters such as co-ordination and muscle strength (Mandillo et al. 2014). Disorders of the central nervous system are often accompanied by deficits in motor function, and affect a number of aspects of movement, from locomotion and balance to finer tasks such as reaching and grasping (Tucci et al., 2007; Preisig et al., 2016). Climbing on the cage and locomotor activity on the cage floor have been found to be important indicators of motor function and form the natural activity routine of mouse motor behavior (Nevison et al., 1999; Borbélyová et al., 2019).

Currently a number of tests, including grip strength and gait analysis (Tucci et al. 2007; Preisig et al. 2016), are used to study the progression of degenerative diseases. These are limited to measuring aspects of motor function, which can also be dependent on external factors, such as experimenter expertise, timing of the test, testing conditions and the motivation of the test subject (Balzani et al., 2018; Baran et al., 2022). In addition, a number of these tests are known to be affected by subtle factors, such as the order of testing and the amount of handling before testing, and crucially, repeated testing in progressive conditions may itself alter the results of subsequent tests (McIlwain et al. 2001; Paylor et al. 2006; Mingrone et al. 2020). Therefore, issues of reproducibility and consistency have to be overcome as researchers strive for greater translatability in preclinical research. This is particularly true for pharmacological investigations that require chronic administration of substances, especially as the short periods of time and potential external confounders affect the integrity and completeness of the result. The probability of clinical success for substances tested in such studies is therefore reduced (Kaffman et al. 2019).

Investigating perturbations in the home-cage activity of undisturbed mice over extended periods can greatly enrich standard out-of-cage phenotyping and provide novel insights into subtle and progressive conditions at early time points. A number of systems have been developed to investigate motor activity over extended periods of time in single-housed as well as group-housed mice. However, measuring both cage-lid climbing and cage-floor movement simultaneously in group-housed mice remained a technical challenge (Bains et al. 2018).

Over the past few years there has been a concerted effort toward overcoming these challenges by housing the mice in testing chambers for extended periods of time and automatically measuring non-evoked activity (Bains et al. 2018). Voluntary wheel running has proved to be a robust indicator of motor-function deficits from an early stage, as it measures a number of motor parameters over several weeks (Lana-Elola et al. 2021). However, concerns such as single housing and the detrimental effect of exercise on certain genetically altered mouse strains that serve as models for diseases such as Huntington’s disease (HD) (Corrochano et al. 2018), including the model used in the current study, in which wheel running may itself affect the phenotype expression, remain an issue. In addition, the subtler indicators of changes in motor function, such as activity around anticipation to light phase change, are not identified through wheel-running activity (Bains et al. 2016).

Despite some early promise shown by gene-targeting therapies, there is currently no disease-modifying treatment for HD, and therapy is focused on management of symptoms and improving quality of life (Kim et al. 2021; Kwon 2021). Progressive loss of neurons is a characteristic feature of this neurodegenerative condition. HD is caused by an unstable expansion within the trinucleotide poly(CAG) tract in exon 1 of the huntingtin gene, located on the short arm of Chromosome 4 (Menalled et al. 2012; Corrochano et al. 2014; Cepeda and Tong 2018), and is characterized by progressive motor deficits such as loss of coordination, tremors and hypokinesis (Schilling et al. 1999). The hallmark histopathology of HD is cell death in the striatum and cerebral cortex, which results in miscommunication between the basal ganglia and the cerebral cortex. This manifests as uncontrolled, involuntary movements (chorea), cognitive deficits and psychiatric symptoms (Cepeda and Tong 2018). The condition is a progressive, ultimately fatal, neurodegenerative disorder.

This study used the B6-TgN(HD82Gln)81Dbo/H, also known as the N171-82Q, model of HD, first published in 1999, in which damage to the basal ganglia structures causes a hyperkinetic disorder (chorea) in combination with a loss of voluntary movements (bradykinesia and rigidity). These phenotypes become evident at around 10.5 weeks of age and manifest as abnormal gait and other behavioral and physiological abnormalities, such as lower grip strength, disturbed limb dynamics and rigidity of the trunk, as well as a tendency toward a lower body weight (Schilling et al. 1999; Ferrante 2009; Preisig et al. 2016). Automated analysis of gait in the lateral and ventral plane has proved to be very useful in the early detection of the subtle changes in limb movement that recapitulate the hyperkinetic phenotype observed in this model, which is detected as early as 10.5 weeks of age (Preisig et al. 2016).

Voluntary locomotion in mouse disease models is highly clinically relevant because it provides an insight into the physiology of the condition, as well as the behavioral motivation of the individual, and is a fundamental readout of the phenotype used in the diagnosis in human patients (Kieburtz et al., 1996; Reilmann et al., 2014). A method that measures the ways in which animals move in non-provoked situations could potentially be a powerful tool for detecting early, and complex, temporal phenotypes.

Advanced image analysis, which can highlight changes in the animal’s gait in both the lateral and the ventral plane, has thus far proven to be the most sophisticated way of extracting subtle motor phenotypes earlier than 13 weeks of age in the model used in our study (Preisig et al. 2016).

In this study, we demonstrate a new tool for the automated analysis of motor activity, which encompasses climbing as well as locomotion on the cage floor, in undisturbed mice over multiple light:dark cycles. Through its application to the N171-82Q model of HD, we have uncovered a robust complex phenotypic profile for disease progression, including early features of motor dysfunction that are fundamental in developing reproducible digital biomarkers for therapeutic testing, especially when targeting the prodromal stages of HD.

In meeting these challenges, we developed an algorithm to automatically annotate climbing behavior from high-definition video captured from a side-on view of the home-cage. The resulting automated climbing behavior annotations provide an important additional parameter set that further enriches the activity profile captured by the Home Cage Analyser system (HCA; Actual Analytics Ltd., UK) (Bains et al. 2016). Our study shows that it is possible to measure two robust indicators of activity simultaneously in group-housed mice from the inbred strain C57BL/6J. Using this technology, we have also demonstrated the advantages of using more comprehensive recording of motor activity to reveal early signs of degeneration in a genetically altered mouse model of HD (N171-82Q (Schilling et al. 1999)).

## 2. Materials and Methods

### 2.1 Animals and Husbandry

All mice used in the study were bred in the Mary Lyon Centre at MRC Harwell and were housed in individually ventilated cages (IVCs; Tecniplast BlueLine 1284) in groups of three mice per cage on Eco-pure spruce chips grade 6 bedding (Datesand, UK), with shredded paper shaving nesting material and small cardboard play tunnels for enrichment. The mice were kept under controlled light (light 07:00–19:00; dark 19:00–07:00), temperature (22 °C ± 2 °C) and humidity (55% ± 10%) conditions. They had free access to water (25 p.p.m. chlorine) and were fed *ad libitum* on a commercial diet (SDS Rat and Mouse No.3 Breeding diet (RM3). All procedures and animal studies were carried out in accordance with the Animals (Scientific Procedures) Act 1986, UK, Amendment Regulations 2012 (SI 4 2012/3039).

For the first study, 18 male and 18 female mice, in six cages of three mice each, from the inbred strain C57BL/6J were recorded at 3 time points: 13–14 weeks, 30–31 weeks and 52–53 weeks of age. For the second study, mice from the mutant strain B6-TgN(HD82Gln)81Dbo/H (HD), were recorded at 3 time points: 8 weeks, 13 weeks and 15–16 weeks of age. Twenty-seven hemizygous (Hemi) males carrying the HD transgene, along with 24 male wild type (WT) littermate controls, and 33 hemizygous females carrying the HD transgene, along with 24 female WT mice, were housed in same-genotype groups of 3 mice per cage. Using the cage as the experimental unit, a sample number of six was calculated to be the most appropriate sample size based on data from previous studies (Supplementary Data). Additional animals were included in the HD study to compensate for the high attrition rate experienced with this model. Therefore, 9 to 11 cages of hemizygous mice were included in the study to avoid under-powering the later time points. Data from all cages were included in the analysis and appropriate statistical methods (described below) were used to account for the differences in the group sizes. Mice were housed with colony-mates born within the same week into cages containing animals of the same genotype.

Three days prior to recording sessions, the animals were transferred to clean home cages with fresh bedding, nesting material and a cardboard rodent tunnel as enrichment material, in line with the standard husbandry procedures for IVC cages. The cages were then placed in an IVC rack in the experimental room for the animals to acclimatize. For each recording, the cages were randomly assigned to an HCA rig. On the first day of recording, each cage was placed onto the ventilation system, within the rig, as would occur during a normal husbandry procedure.

Animal welfare checks were carried out visually twice daily. At the end of the recording period, the home cages were removed from the HCA rigs and returned to their original positions on the IVC racks.

### 2.2 Microchipping

Radio frequency identification microchips were injected subcutaneously into the lower left or right quadrant of the abdomen of each mouse at 12 weeks of age for the C57BL/6J study and 7 weeks of age for the B6-TgN(HD82Gln)81Dbo/H study. These microchips were contained in standard ISO-biocompatible glass capsules (11.5 × 2mm; PeddyMark Ltd., UK). The procedure was performed on sedated mice (Isoflo; Abbott, UK) after topical application of local anesthetic cream on the injection site prior to the procedure (EMLA Cream 5%; AstraZeneca, UK). The animals were allowed to recover from the microchip procedure for at least one week before being placed in the HCA rigs for data collection. The procedure has been described previously in Bains et al., (2016).

### 2.3 Measurement of Climbing and Validation

Climbing behavior is measured on a frame-by-frame basis by numerically characterizing the pattern of motion occurring within a pre-defined region around the cage lid and quantifying its similarity to a set of key reference examples of climbing and non-climbing behavior (selected programmatically from a large set of human “training” annotations) to yield a classification decision. More specifically, the local trinary pattern representation proposed in Yeffet and Wolf (2009), is used to characterize motion within a 690 × 385 pixel region adjacent to the cage lid as a 16384-dimensional vector; this particular representation was shown in Burgos-Artizzu et al. (2012) to provide an effective, yet computationally efficient, means of distinguishing between different mouse behaviors. Local trinary pattern vectors were extracted for every video frame across more than 7 hours of human-annotated video footage—encompassing over 130 separate bouts of climbing—and were used to train a linear support vector machine classifier (SVM) (Fan et al. 2008) to distinguish between climbing and non-climbing instances. To leverage the correlation between consecutive video frames, a temporal voting window was applied, such that the final classification of a given video frame represented the consensus over a wider time period spanning the frame in question (the underlying logic is to reduce spurious “single frame” detections, while conversely preventing erroneous “splitting” of longer bouts of climbing on the basis of a single misclassified frame.) A leave-one-out cross-validation procedure was applied over the available “training” set of 15 discrete 30-minute video segments, in order to identify: i) the SVM regularization parameter values and ii) the temporal aggregation parameters that—on average—yielded the best generalization performance. The final classifier, generated from the full set of available training data using the parameters revealed by the preceding cross-validation process, was then tested on a further 2.5 hours of — unseen — annotated test videos, yielding 85.6% frame-by-frame accuracy (where 65.9% of climbing frames, and 94.3% of non-climbing frames were correctly classified, with climbing behavior accounting for approximately 30% of the test data). Considering this test data in terms of 5 minute time bins, automatically annotated climbing time correlates well with human annotated climbing time, as confirmed by Spearman’s rank coefficient (ρ=0.836; n=30; p<0.00000001).

### 2.4 Data Analysis

#### 2.4.1 Linear mixed-effects modeling

To account for dependence between data recorded over separate days from the same cage (repeated measures) and to avoid pseudo-replication, statistical analyses were conducted using linear mixed-effects modeling. To adjust for parameters with non-normal distributions, data were box-cox transformed prior to analysis. Any subsequent modeling satisfied assumptions of normally distributed residuals.

We constructed a linear mixed-effects model of either distance moved or time spent climbing (continuous variables) as a function of the effect of sex, age of caged mice and genotype (all categorical fixed effects). Cage ID was modeled as the random effect intercept with day of recording as the random effect slope. This structure allows for cages to vary randomly in their baseline distance moved or time spent climbing value, and for this relationship to vary randomly according to day of recording. It will account for time-dependent and cage-specific fluctuations in activity over the three days of recording.

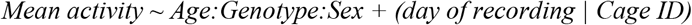

This model was compared to other model iterations with different combinations of sex, age and genotype with or without the interaction term and random effects structure. An ANOVA test was run to determine the statistical significance of the interaction between Age:Genotype:Sex and to inform model selection. Random effects and fixed effects found not to have a statistically significant contribution to model fit were eliminated. Models were fit using R’s “lmer” function.

#### 2.4.2 Timeframes of interest

The timeframes of interest in the current study were defined as the 30 minutes directly preceding lights being turned on (06:30 to 07:00) and 30 minutes directly preceding lights being turned off (18:30 to 19:00). This definition was consistent between both parameters of interest: distance moved (mm) and time spent climbing (seconds). When analyzing activity during the timeframes of interest, we first summed data per time bin (6 min) for mouse within a cage. We then calculated the average activity per cage across the 5 time bins.

We have previously shown that C57BL/6J mice show peak activity in the dark phase, and that mouse activity varies around change in light phases in a strain-specific manner (Bains et al. 2016). In the current study these changes were particularly relevant, as mouse models of neurodegenerative diseases, including HD, have known sleep disturbances. Sleep onset latency towards the end of the active period is a particularly sensitive measure (Morton et al. 2005), therefore the first 30 minutes and the last 30 minutes of the active period were chosen as the timeframes of interest for further investigation.

#### 2.4.3 Post-hoc analysis

We conducted pairwise post-hoc comparison tests by computing the estimated marginal means (least-squares means) for factor combinations and correcting for multiple comparisons using the Benjamini–Hochberg method to decrease the false discovery rate. This process was run using R’s “emmeans” function, which returned adjusted p values. These values were used to indicate the statistical significance of the genotype effect at various levels of factor combination.

While the algorithm for automated behavior annotation is proprietary, the analysis is openly available as a part of this manuscript; please see supplementary material. The datasets for the experiments in this manuscript are also openly available on request.

All climbing data were converted from number of frames to time spent climbing in seconds prior to analysis and visualization, as the authors believe that to be a more intuitive parameter. The number of frames is converted to time in seconds as follows:

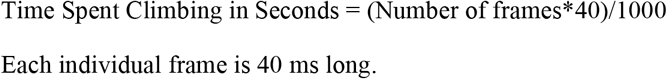

## 3. Results

### 3.1 Multiday recording in mouse home cage shows sexual dimorphism in C57BL/6J mice and reveals age-related decrease in activity

The data from the females of inbred strain C57BL/6J show significantly higher cage-floor activity, described as distance traveled in mm (Fig. 1G) in the dark phase, as compared to males at 3 months of age (n = 6, p < 0.0001). This difference in cage floor activity during the dark phase persists as the mice age, as seen in the later time points of 7 months (p = 0.0001) and 12 months (p < 0.01) (Fig. 1G, n = 6). The activity shows a clear circadian rhythm (Fig. 1).

**Figure 1.**
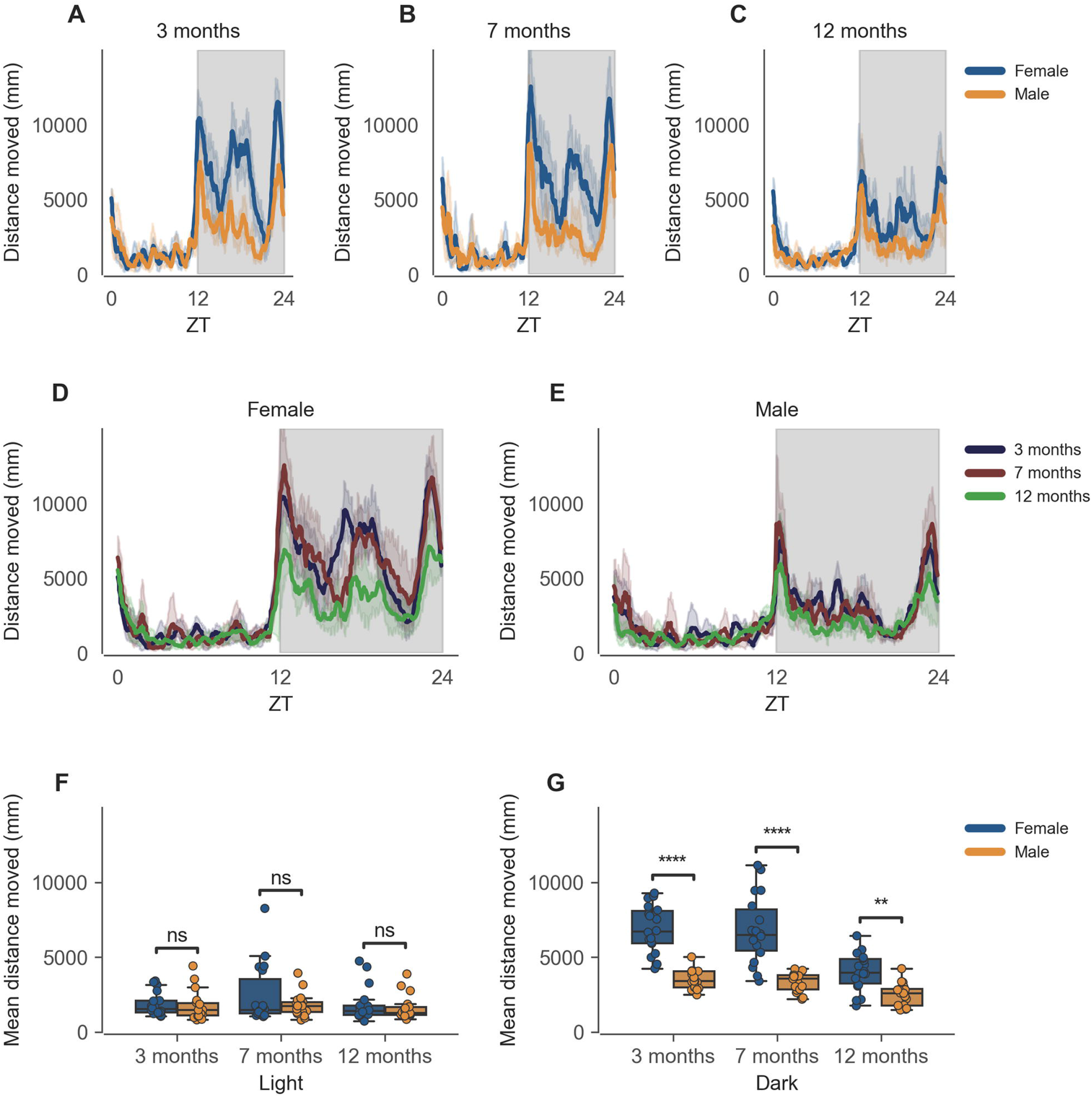
Distance moved by mice during dark phase varies according to sex and across age during passive home-cage monitoring. **(A)** Distance moved (mm) over zeitgeber time in female and male cages of 3-month-old mice during recording session, binned into 6-min time bins and averaged over a 24-hour period. Line represents mean distance over time across cages of a sex group; shaded error band represents 95% confidence interval. Data from individual mice within a cage were summed to produce one time series per cage. Grey shaded areas on plot represent darkness. **(B, C)** Same as **A** but for 7-month-old and 12-month-old mice, respectively. **(D)** Distance moved over zeitgeber time in female cages of mice of all ages. Line represents mean distance over time across cages of a sex and age group; error-shaded area represents 95% confidence interval. **(E)** Same as **D** but for male cages. **(F)** Boxplot of mean distance moved during light phase with cages split by age and by sex. Distance moved within light phase was averaged across time per cage and per day. Three data points per cage (three days of recording) were modelled using a linear mixed-effects model to account for repeated-measures and least-squares means estimated to return adjusted p values of levels of factor combinations. **(G)** Same as D but for dark phase. **p<0.05, ****p<0.0001

### 3.2 Automated climbing annotation in mouse home cage records complex sexual dimorphism in C57BL/6J mice and reveals age-related decrease in activity

The data from the females of inbred strain C57BL/6J show significantly higher cage-bar climbing, described as time spent climbing in seconds (Figure 2), as compared to males at 3 months of age (Fig. 2G, n = 6, p < 0.0001), this difference in cage-bar climbing during the dark phase persists as the mice age as seen at the later time points of 7 months (Fig. 2G, n = 6, p < 0.0001) and 12 months (Figure 2G, n = 6, p < 0.0001). The activity shows a clear circadian rhythm (Fig. 2).

**Figure 2.**
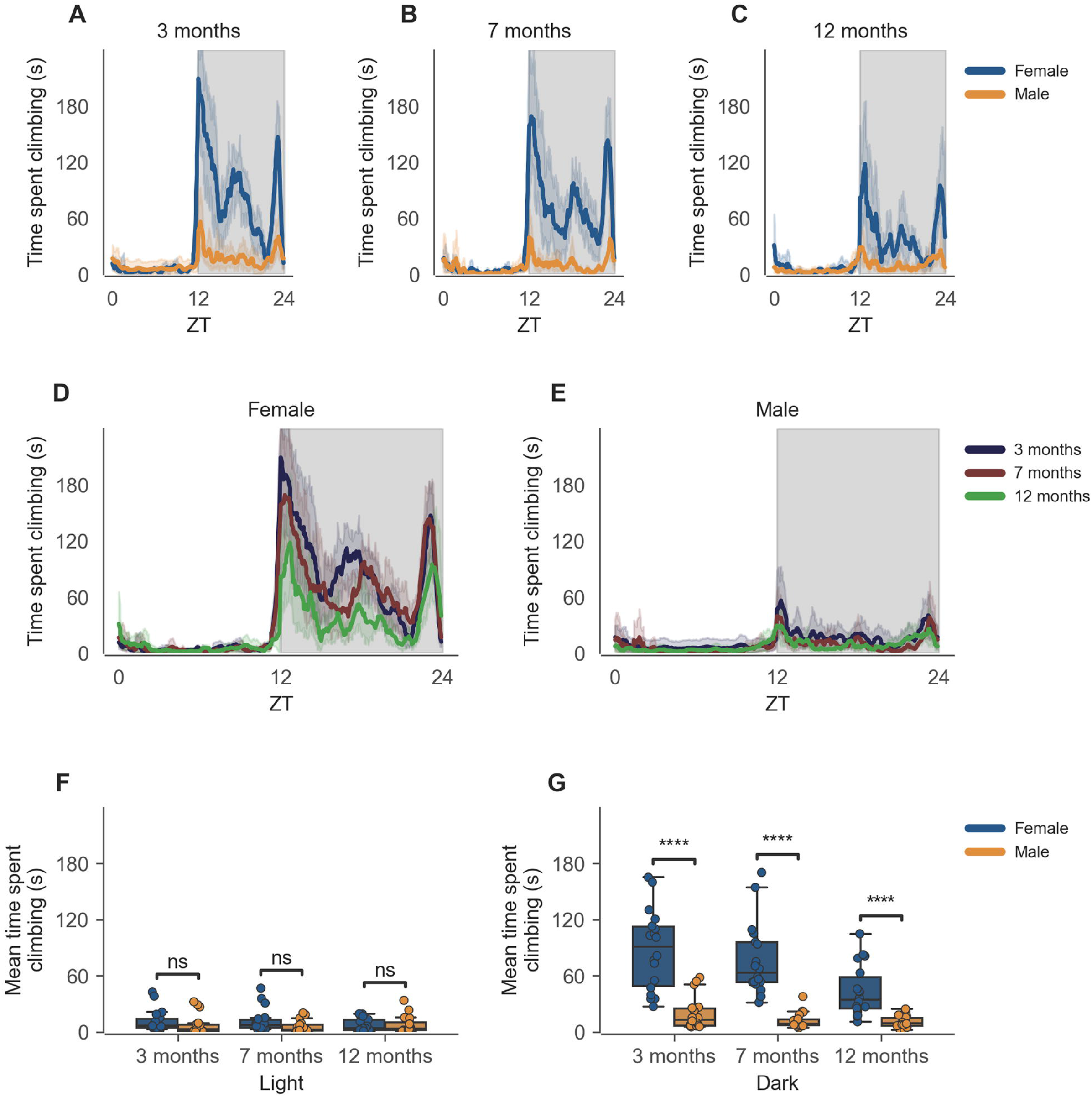
Time spent climbing by mice during dark phase varies according to sex and across age during passive home-cage monitoring. **(A)** Time spent climbing over zeitgeber time in female and male cages of 3-month-old mice during recording session, binned into 6-min time bins and averaged over a 24-hour period. Line represents mean time spent climbing over time across cages of a sex group; shaded error band represents 95% confidence interval. Data from individual mice within a cage were summed to produce one time series per cage. Grey shaded areas on plot represent darkness. **(B, C)** Same as **A** but for 7-month-old and 12-month-old mice, respectively. **(D)** Time spent climbing over zeitgeber time in female cages of mice of all ages. Line represents mean distance over time across cages of a sex and age group; error-shaded area represents 95% confidence interval. **(E)** Same as **D** but for male cages. **(F)** Boxplot of mean time spent climbing moved during light phase with cages split by age and by sex. Time spent climbing within light phase was averaged across time per cage and per day. Three data points per cage (three days of recording) were modelled using a linear mixed-effects model to account for repeated-measures and least-squares means estimated to return adjusted p values of levels of factor combinations. **(G)** Same as D but for dark phase. ****p<0.0001

### 3.3 Early detection of activity phenotype in mouse model of Huntington’s disease

The data from the HD model recapitulate the clear circadian rhythm and sex differences seen in the inbred strain, in which females were significantly more active than males in both measures of activity. The total activity over the dark and light phases of hemizygous (Hemi) HD mice compared to the WT HD mice was not statistically different from each other for both sexes.

There is, however, a specific time-of-day-dependent deficit in activity seen in Hemi HD mice as compared to WT HD mice, at the transition between dark and light phases for females (06:30 to 07:00). This difference was apparent as early as 8 weeks, when the animals showed no overt signs of the disease onset (Fig. 3E, n = 8 WT/11 Hemi, p < 0.01). The decrease in cage-floor activity at this time became even more apparent at 13 weeks of age (Fig. 3E, n = 7 WT/7 Hemi, p < 0.0001). By 15–16 weeks of age the mice showed clear signs of disease and differences between the two genotypes were distinct (Fig. 3E, n = 7 WT/3 Hemi, p < 0.0001).

**Figure 3.**
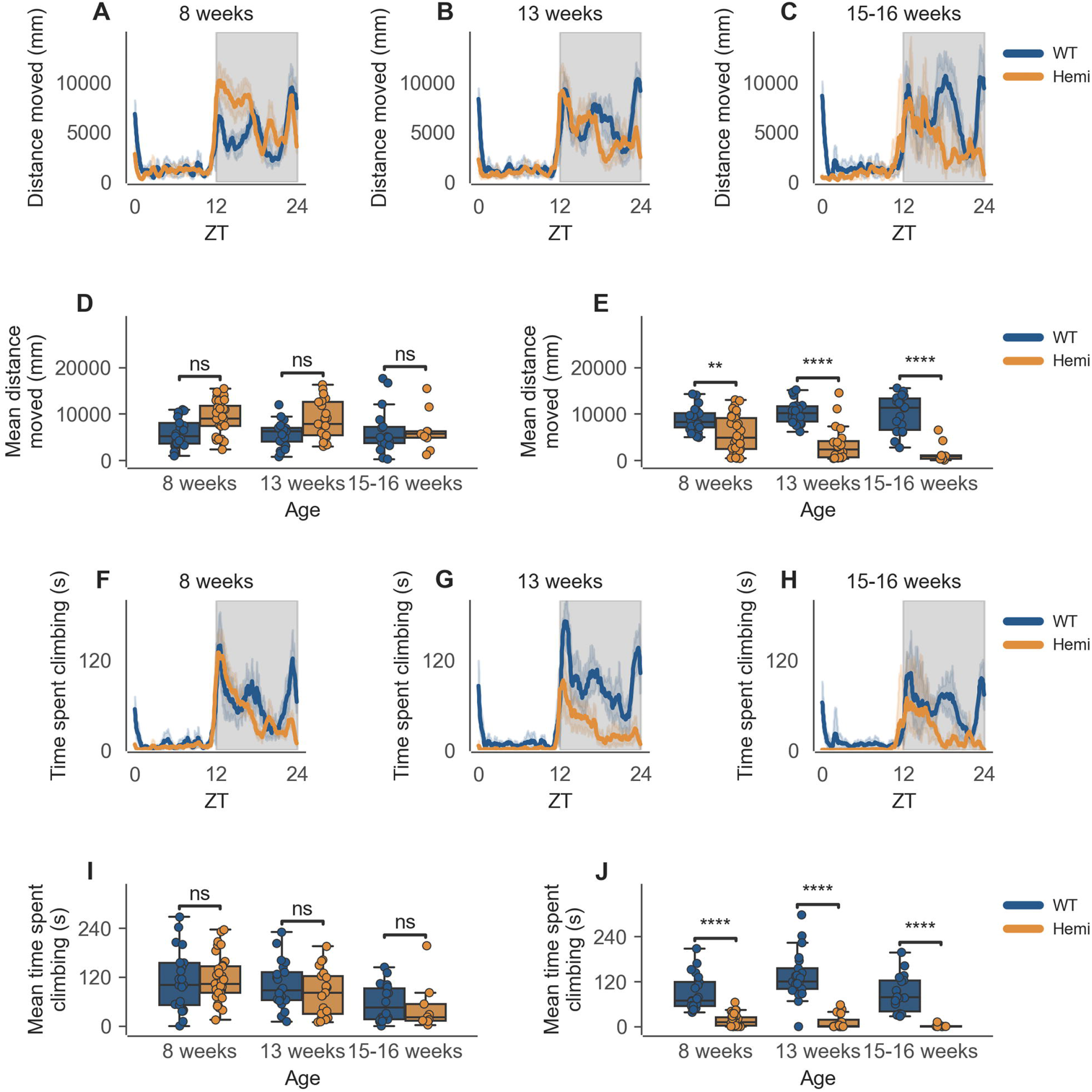
Distance and climbing activity varies according to genotype in female mice and across age at conclusion of dark phase, but not at beginning of dark phase. **(A)** Distance moved over zeitgeber time in female-only cages of 8-week-old mice, split according to genotype, during recording session, binned into 6-min time bins and averaged over a 24-hour period. Line represents mean distance over time across cages of a genotype group, shaded error band represents 95% confidence interval. Data from individual mice within a cage was summed to produce one time-series per cage. Grey shaded areas on plot represent darkness. **(B, C)** Same as **A** but for 13-week-old and 15-16-week-old mice, respectively. **(D)** Boxplot of mean distance moved during first 30 minutes of darkness within female-only cages split by age and genotype. Distance moved was averaged across first 30 minutes of darkness per cage and per day. Three data points per cage (three days of recording) were modelled using a linear mixed-effects model to account for repeated-measures and least-squares means estimated to return adjusted p values of levels of factor combinations. **(E)** Same as **D** but for last 30 minutes of darkness. **(F)** Time spent climbing over zeitgeber time in female-only cages of 8-week-old mice, split according to genotype, during recording session, binned into 6-min time bins and averaged over a 24-hour period. Line represents mean time spent climbing over time across cages of a genotype group, shaded error band represents 95% confidence interval. Data from individual mice within a cage was summed to produce one time-series per cage. Grey shaded areas on plot represent darkness. **(G, H)** Same as **F** but for 13-week-old and 15-16-week-old mice, respectively. **(I)** Boxplot of mean time spent climbing during first 30 minutes of darkness within female-only cages split by age and genotype. Time spent climbing was averaged across first 30 minutes of darkness per cage and per day. Three data points per cage (three days of recording) were modelled using a linear mixed-effects model to account for repeated-measures and least-squares means estimated to return adjusted p values of levels of factor combinations. **(J)** Same as **I** but for last 30 minutes of darkness. **p<0.01, ****p<0.0001

This decrease in activity at the end of the dark phase (06:30 to 07:00) is also seen in male Hemi HD mice as compared to WT HD mice, allowing for the phenotype to be detected as early as 8 weeks of age (Fig. 4E, n = 8 WT/9 Hemi, p < 0.05), persisting into the next time point of 13 weeks of age (Fig. 4E, n = 8 WT/6 Hemi, p < 0.05), and becoming obvious at 15–16weeks of age (Fig. 4E, n = 8 WT/5 Hemi, p < 0.001).

**Figure 4.**
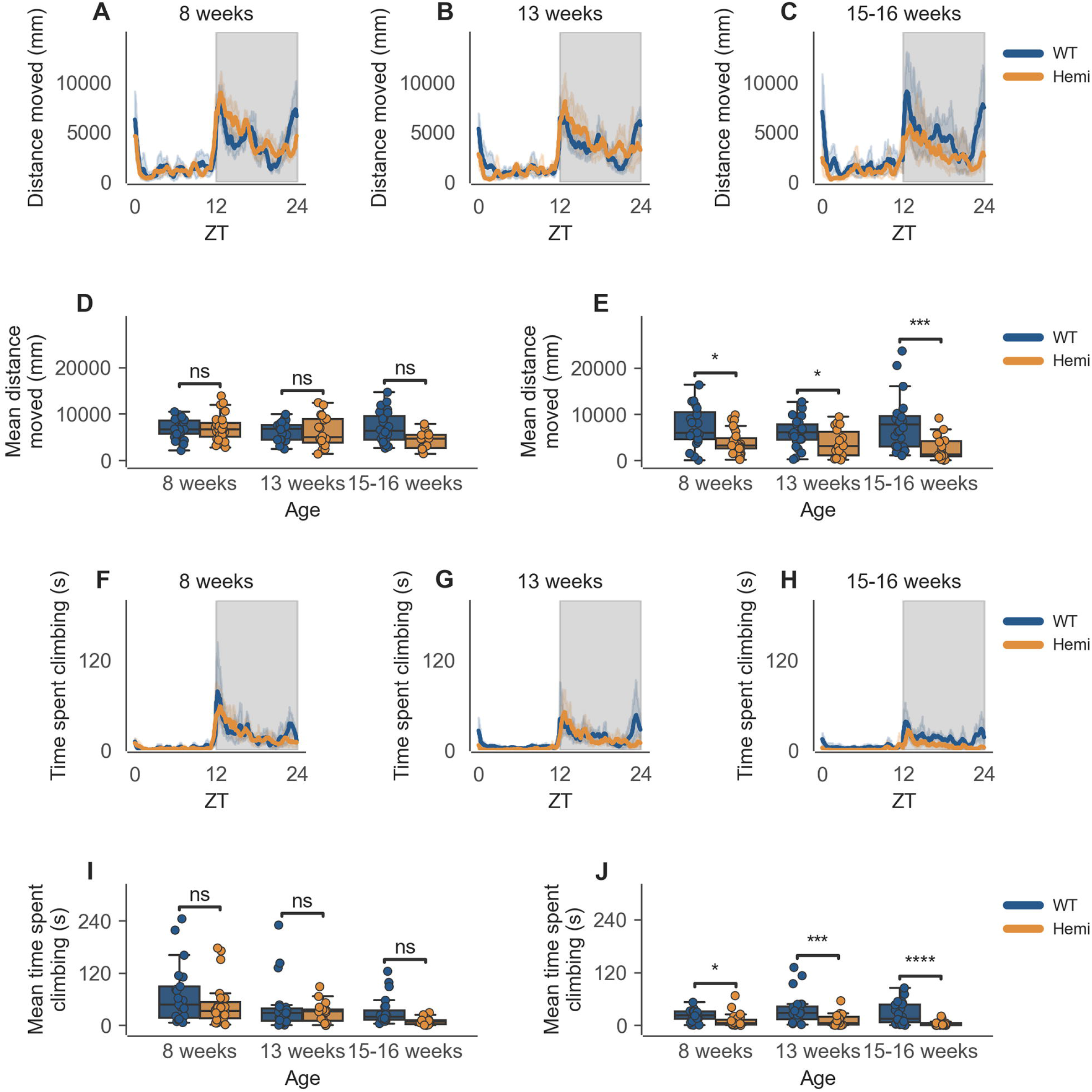
Distance and climbing activity varies according to genotype in male mice and across age at conclusion of dark phase, but not at beginning of dark phase. **(A)** Distance moved over zeitgeber time in male-only cages of 8-week-old mice, split according to genotype, during recording session, binned into 6-min time bins and averaged over a 24-hour period. Line represents mean distance over time across cages of a genotype group, shaded error band represents 95% confidence interval. Data from individual mice within a cage was summed to produce one time-series per cage. Grey shaded areas on plot represent darkness. **(B, C)** Same as **A** but for 13-week-old and 15-16-week-old mice, respectively. **(D)** Boxplot of mean distance moved during first 30 minutes of darkness within male-only cages split by age and genotype. Distance moved was averaged across first 30 minutes of darkness per cage and per day. Three data points per cage (three days of recording) were modelled using a linear mixed-effects model to account for repeated-measures and least-squares means estimated to return adjusted p values of levels of factor combinations. **(E)** Same as **D** but for last 30 minutes of darkness. **(F)** Time spent climbing over zeitgeber time in male-only cages of 8-week-old mice, split according to genotype, during recording session, binned into 6-min time bins and averaged over a 24-hour period. Line represents mean time spent climbing over time across cages of a genotype group, shaded error band represents 95% confidence interval. Data from individual mice within a cage was summed to produce one time-series per cage. Grey shaded areas on plot represent darkness. **(G, H)** Same as **F** but for 13-week-old and 15-16-week-old mice, respectively. **(I)** Boxplot of mean time spent climbing during first 30 minutes of darkness within male-only cages split by age and genotype. Time spent climbing was averaged across first 30 minutes of darkness per cage and per day. Three data points per cage (three days of recording) were modelled using a linear mixed-effects model to account for repeated-measures and least-squares means estimated to return adjusted p values of levels of factor combinations. **(J)** Same as **I** but for last 30 minutes of darkness. *p<0.05, ****p<0.0001

This specific time-of-day-dependent deficit in cage-floor activity is mirrored in cage-bar climbing, described as time spent climbing, where female Hemi HD mice show a significant decrease in time spent climbing compared to female WT HD mice, at the transition between dark and light phases (06:30 to 07:00). Once again, this difference was apparent as early as 8 weeks, when the animals showed no overt signs of the disease onset (Fig. 3J n = 8 WT/11 Hemi, p < 0.0001). As seen with cage-floor activity, the decrease in cage-bar climbing at this time became even more apparent at 13 weeks of age (Fig. 3J, n = 7 WT/7 Hemi, p < 0.0001) and by 15–16 weeks of age the mice showed clear signs of disease and differences between the two genotypes were distinct (Fig. 3J, n = 7 WT/3 Hemi, p < 0.0001).

As with cage floor activity, time spent climbing also follows the same pattern in male Hemi HD mice as compared to male WT HD mice, where a statistically significant difference in genotypes is observed at 8 weeks of age (Fig. 4J, n = 8 WT/9 Hemi, p < 0.05), persisting to the next time point of 13 weeks of age (Fig. 4J, n = 8 WT/6 Hemi, p < 0.001) and becoming obvious at 15–16weeks of age (Fig. 4J, n = 8 WT/5 Hemi, p = 0.0001).

## 4 Discussion

Classically, with a few exceptions, all behavior testing is carried out during the light phase, in which resting animals are removed from their home cage and placed in a novel environment away from their cage mates (Bains et al. 2018). Such out-of-cage tests are known to be influenced by factors such as ambient noise, lighting and odors, and handling methods, resulting in stress and anxiety-like responses. Home-cage monitoring, by contrast, is free of such influences as these behavioral confounds are removed. Therefore, the welfare burden on the animals is much lower (Voikar and Gaburro 2020) as behaviors can be recorded outside of the normal observation hours, allowing for continual monitoring of progressive phenotypes over both light and dark cycles and not just at specific time points. Such a one-size-fits-all approach risks missing critical milestones for disease emergence and/or progression. This is particularly so as the home-cage activity declines with age (Nakamura et al. 2011; Yanai and Endo 2021) and snapshots of these milestones may not be enough to reveal complex phenotypes that change with time of day, or are particularly exaggerated at certain times of the day in relation to the light:dark cycle, as shown in this study.

The advantage of observing the mice undisturbed within their home cage over multiple light–dark cycles is that, in addition to the observed phenotype, it is also possible to disentangle the temporal appearance of such phenotypic traits. The data from C57BL/6J show that there is a clear sexual dimorphism in the overall activity of the animals, and that both males and females show peak activity in the dark phase. These data therefore point to a clear circadian influence. This method of analysis allows one to also investigate the influence of ultradian parameters on measures such as activity and climbing. The importance of this finding is highlighted in the HD study, in which the males show very low baseline activity in both mutant and WT strains. We have already shown that the HCA system is capable of detecting statistically significant changes in activity around the light phase changes between various background strains (Bains et al. 2016). Here we extend this concept to draw out clinically relevant, subtle and early phenotypic changes by focusing on specific times of interest such as the first 30 and last 30 minutes of the dark phase.

Through the current study, we showcase a recently developed automated behavior annotation tool for climbing behavior in standard IVCs under group-housed conditions. We have previously shown the capabilities of the system in investigating cage-floor activity in group-housed mice in their home cages (Bains et al. 2016). Here, we show that measuring climbing is part of the standard motor behavior repertoire of mice and can greatly enhance the existing dataset to investigate motor phenotypes in much greater detail and with minimal experimenter intervention.

Sexual dimorphism in climbing behavior has been reported previously in singly tested C57BL/6Ntac mice, using the LABORAS system, in which the main aim of the study was to investigate the difference in response to novelty between the sexes. Furthermore, parameters were only measured for 10 minutes (Borbélyová et al. 2019). More recently, a study on the effect of age, sex and strain on cage-lid climbing in single-housed mice has also reported sexual dimorphism, as well as strain and age differences, in single-housed mice, over 24 hours, peaking in the dark phase (Zhang et al. 2021). The data from the inbred strain C57BL/6J, in the current study show significantly high cage-bar climbing as well as cage-floor activity in females as compared to males at all three age time points, with most of the activity observed in the dark phase. To the best of our knowledge, this is the first system of its kind that can detect cage-level climbing activity in group-housed mice within their home cage for extended periods of time, without the need for removal into specialized and/or novel caging.

Serious motor and cognitive deficits that are the hallmark of HD are often preceded, by decades, by more subtle changes in circadian rhythms and motor function (Wang et al. 2018; Wiatr et al. 2018). Therefore, an approach that screens for the chronic and progressive nature of such conditions is more clinically relevant than one that screens for acute signs of motor deficits that manifest at a much later stage of the disease. One such approach is to focus on behaviors that are elective and not essential to survival, such as grooming, playing or climbing, as they reflect the animal’s emotional or motivational state, which would be ethologically more relevant for a preclinical model (Zhang et al. 2021). Therefore, a perturbation in such behaviors could reflect a suboptimal health state, especially in conditions that are chronic and progressive rather than acute.

The onset of HD is often insidious and progressive and the phenotypes are biphasic; at early stages of the condition involuntary functions are affected and in the later stages the directly controlled, voluntary functions begin to fade. This means that motor phenotypes are often expressed as hyperkinesia in the early stages and akinesia in the later stage (Kim et al. 2021). Therefore, it becomes imperative to investigate such conditions longitudinally and for extended periods of time as the ‘snapshot in time’ investigations, such as those that investigate motor activity in an open field, may not be representative of a clinically relevant disease profile.

The HD data in the current study show a significant increase in signal in both cage-floor and cage-lid climbing activities around the transition between the light-to-dark and dark-to-light phases. There is ample evidence to show that spontaneous cage-bar climbing is mediated through the dopaminergic system and therefore depends on the motivation and arousal state of the mouse (Joshua et al. 1982; Palmiter 2008; Brooks and Dunnett 2009). As mice are active during the dark phase, the arousal states coincide with the transition periods between light and dark phases of the circadian cycle; we have already shown that the most active periods as seen from cage-floor activity, are around these transition times (Bains et al. 2016). The current study shows that this is also true for cage-bar climbing. In addition, despite the decrease in total activity with age, this increase in cage-bar climbing and cage-floor activity around dark-to-light phase transition persists. This aspect is of particular interest in those models in which the genetic manipulation modeling the disease would result in a greater decrease in activity with age, as compared to wild-type counterparts. However, as the activity in the wild-type mice also decreases with age, any decrease in activity due to the genotype is therefore hard to discern in conventional testing paradigms.

The importance of this finding is highlighted in the set of experiments carried out using the mouse model of HD, N171-82Q. These data recapitulate the sex differences seen in the C57BL/6J strain experiments, in which females were significantly more active than males in both measures of activity, with this difference persisting across all time points. However, of note is a specific time-dependent decrease in cage-bar climbing activity at the transition between dark and light phases, which was apparent as early as eight weeks, when the animals showed no overt signs of the disease onset. The decrease in cage-floor activity at this time was also observed at 8 weeks of age; however, the decrease in cage-floor activity became even more apparent at 13 weeks of age. By 15–16 weeks of age the mice showed clear signs of disease and differences between the two genotypes were distinct. This last finding is of particular interest as clinical case studies, as well as mouse models of HD, are known to present with sleep disturbances, one of the hallmarks of the condition (Pallier et al. 2007). Whilst the mechanism is not fully understood, there is some evidence that this change may be attributed to increased pathology either in the brain region controlling circadian rhythms—the suprachiasmatic nucleus—or in a pathway further downstream (Pallier et al. 2007).

This study recapitulates the aspect of the disease in which the offset of activity in HD mice is observed sooner than that in their WT counterparts, for both cage-floor activity, as well as cage-bar climbing. Indeed, a 2005 study comparing the circadian activity patterns of human patients with a different mouse model of Huntington’s disease (R6/2), reported a similar pattern of decline in activity towards the end of the active phase with disease progression (Morton et al. 2005). In human patients, this manifests as spending a longer time in bed. In the absence of a complete circadian screen, which would be outside the scope of this study, it would not be over-anthropomorphizing to say that HD mice begin their rest period earlier than their wild-type counter parts and remain at rest for longer, from the earliest stages of the disease.

In progressive degenerative conditions, neuronal dysfunction occurs before any overt signs of the condition become apparent in the behavior. As neurons are unable to regenerate, most therapies under development focus on neuroprotection, with the aim of slowing the progression of the disease and, where possible, delaying the onset (Jin et al. 2014; Kumar et al. 2015). This necessitates the development of models in which the therapeutic window aims to target the pre-manifestation period, in order to minimize neuronal loss. Therefore, any models that can identify the earliest manifestation of the mutation are invaluable in investigation of the disease progression and identification of early biomarkers (Levine et al. 2004).

It is important to remember, however, that animal behavior is complex, and that external factors, such as effects of diet and exercise, can have an impact on disease progression (Dutta et al. 2021). The model used in the current study, N171-82Q, is known to have a more variable phenotype than that of the more severe R6/2 model, even though the motor phenotype and weight loss generally becomes evident at 11 weeks of age (Ferrante 2009). Such disease models are generally complex in their development, and part of the required improvement in animal research is the development of tools with the ability to capture this complexity both in terms of different phenotypes measured and the timings of their appearance. The factors driving spontaneous cage-lid climbing are not fully understood, but it is clear that this activity is affected by a decline in welfare (Zhang et al. 2021). Thus, we can say that investigating the non-evoked total motor function repertoire of animals in progressive and degenerative conditions is the first step toward early phenotype recognition and can be extended to other mutant models showing complex phenotypes.

## 6 Author Contributions

RB was responsible for the experimental design, experimental procedure, data collection and manuscript preparation. HF carried out bioinformatics and statistical analysis. RS was responsible for the system design, including automated climbing algorithm. JA contributed to the study design and system design. MS contributed to the study design. PN contributed to the study design, carried out circadian and activity data analysis, and prepared the manuscript. SW contributed to the study design, including animal procedures, and prepared the manuscript. All authors contributed to the article and approved the submitted version.

## 8 Acknowledgments

We wish to thank the IT Infrastructure team at the Mary Lyon Centre for their support with the hardware. We also wish to thank the animal care team at the Mary Lyon Centre for their help and technical support. A special thanks to Mrs. Louise Tinsley, for proof reading and formatting this manuscript. Finally, we wish to thank the National Centre for 3Rs for their continued support of the home-cage concept.

## 12 Data Availability Statement

The datasets analyzed for this study can be found in the github repository [https://github.com/HamishForrest/Bains-et-al.-2023]. This is a public repository. The raw datasets for the experiments in this manuscript are also openly available on request.

